# Statistical signals indicate a dependence between amino acid backbone conformation and the translated synonymous codon

**DOI:** 10.64898/2026.04.02.712692

**Authors:** A. A. Rosenberg, A. Marx, A. M. Bronstein

## Abstract

Synonymous codons encode the same amino acid but can differ in their usage and translational properties. In previous work we reported statistical differences in backbone dihedral angle distributions associated with synonymous codons in the Escherichia coli proteome. This finding has been questioned due to concerns regarding the statistical methodology used. Here we revisit the dataset using corrected statistical procedures and alternative statistical tests. Across multiple frameworks, the real dataset consistently shows an excess of small p-values relative to randomized controls, indicating detectable codon-associated differences in backbone conformation.

## Main Text

Codon usage bias (CUB) is a widespread evolutionary phenomenon observed across all domains of life. Synonymous codons encode the same amino acid but can differ in frequency and translational properties, reflecting a complex interplay between mutation, selection, and cellular constraints on translation.

In previous work, we investigated whether synonymous codon identity is associated with local protein backbone conformation by comparing Ramachandran angle distributions conditioned on synonymous codons (1). Using a statistical framework based on permutation tests and kernel density estimates, we detected statistically significant differences between the backbone dihedral angle distributions associated with synonymous codons. Importantly, we emphasized that this observation represented a statistical association rather than a demonstrated causal relationship. Several possible explanations were proposed, including differences in translation kinetics between codons or evolutionary selection acting on codon usage in structurally distinct regions.

Concurrent with our report that synonymous codon usage is associated with local protein backbone dihedral angles, Cope and Gilchrist (2) used a population-genetics model (ROC-SEMPPR) to account for gene expression and amino-acid biases, concluded that selection on synonymous codon usage varies only weakly across protein structural regions and that codon-dependent structural signals could largely be explained by global selection for translational efficiency rather than by region-specific associations with structure. A later analysis of our dataset (3) argued that our original statistical procedure may be overly sensitive due to the kernel density estimation parameters used in the test statistic and suggested that the observed codon-dependent structural signals could be explained by gene expression–related selection on codon usage.

More recently, González-Delgado et al. (4) noted a theoretical flaw in our statistical test due to the bootstrapping procedure we used to combine permutation tests over size-matched samples from each pair of codons, showing that the approach is statistically valid only as a single permutation test, without the bootstrap procedure. The authors suggest that the density estimates describing the samples may therefore be biased and invalid. Using an alternative two-sample test of distributional equality not based on permutations, as well as a test statistic measuring the Wasserstein distance between 1-dimensional projections on a torus (5), these authors reported that no statistically significant codon-dependent differences in backbone dihedral angles could be detected.

Given these critiques, it is important to reassess whether any statistical signal linking synonymous codons and backbone conformation remains detectable when using statistically valid procedures and alternative test statistics. Here we revisit the original dataset using an updated analysis framework finding that across all tested statistical frameworks, we consistently observe differences between codon-conditioned backbone angle distributions in the real dataset that are absent in the randomized control. These results indicate that a statistical signal linking synonymous codon identity and backbone conformation remains detectable and is robust to the choice of statistical test. While the biological origin of this signal remains unclear, its persistence across independent statistical methods suggests that further investigation into codon–structure relationships is warranted.

To re-analyze the dataset introduced in our previous work (1), we removed the bootstrap resampling procedure used in our original analysis. Instead, we performed standard permutation tests with an increased number of permutations (K = 5000) to maintain comparable statistical power while ensuring validity of the test. Two classes of test statistics were evaluated:

1. **KDE-L1 statistic** used in the original work (1)
2. **Projected Wasserstein distance on the torus**, as proposed in (5)

In addition, we implemented the full statistical testing framework introduced by González-Delgado et al., which directly computes p-values for Wasserstein-based statistics without permutation testing.

To assess whether the statistical procedures produce spurious rejections under the null hypothesis, we constructed a randomized control dataset where codons were randomly assigned to each amino acid according to empirical codon frequencies measured within the same secondary-structure class across the E. coli proteome (denoted as **Randomized (AA+SS prior)**). Because codon identity is randomized within amino-acid categories, any statistically significant differences detected in this dataset would indicate excessive sensitivity of the statistical test rather than a genuine signal. In all the following analyses, we used the randomized data as the control and removed the tests comparing a codon to itself.

The original KDE-based analysis using the corrected permutation framework without bootstrapping results in the distributions of p-values for synonymous codon comparisons shown in Figure 1A. Across a wide range of kernel bandwidth parameters, the p-value distributions obtained from the real dataset differ markedly from those obtained from the randomized control dataset.

**Figure 1:**
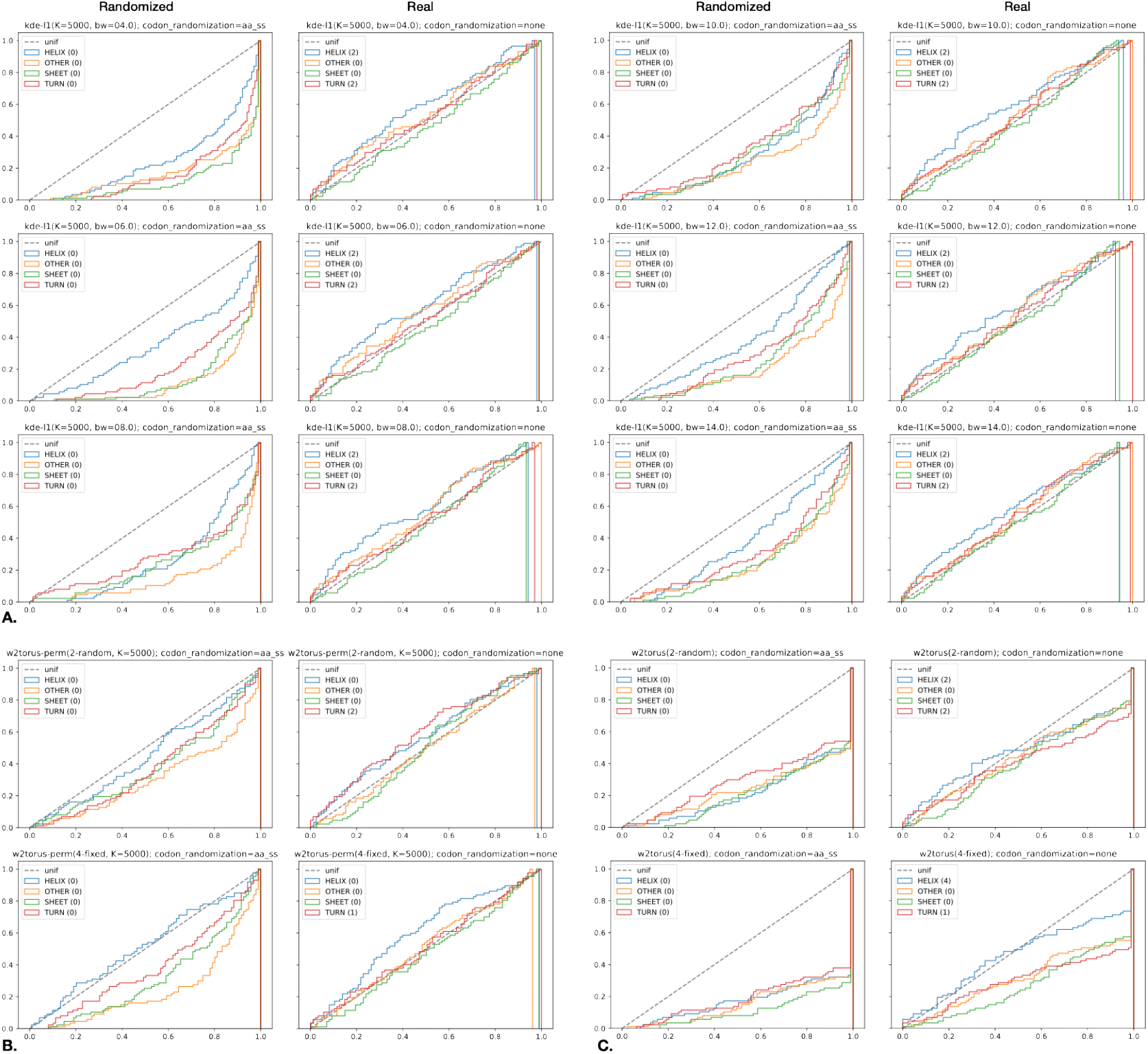
Cumulative distributions of p-values produced by the different tests. (A) Permutation test with KDE-L1 test statistic (1), with different KDE kernel bandwidths; (B) Permutation test with the projected Wasserstein distance test statistic (5), with either 2 or 4 projections, chosen randomly or fixed; and (C) the statistical test proposed by González-Delgado *et al*. (5). All plots are paired, showing distributions on the **Real** and **Randomized (AA+SS)** sets. In all cases the distribution of p-values under the null (randomized set) is super-uniform and it appears consistently and similarly different between the two sets strongly suggesting the existence of a structure-coding signal. Only the **Real** dataset results in rejections of the null hypothesis further suggesting significance of the signal. Number of rejections in each secondary structure category is reported in the legends.

Specifically, p-values derived from the randomized dataset follow a super-uniform distribution, as expected under the null hypothesis. In contrast, the real dataset exhibits a pronounced excess of small p-values, resulting in multiple rejections of the null hypothesis under Benjamini–Hochberg false-discovery-rate control (FDR = 0.05).

To determine whether this signal is an artifact of our specific method (permutation test with KDE-L1 test statistic), we repeated the analysis using the projected Wasserstein distance on the torus proposed by González-Delgado et al. (5), using their original R implementation (6) to compute the test-statistic value. The Wasserstein test statistic requires the choice of projection directions, so we experimented with 2 or 4 projection directions, chosen either randomly or fixed (based on the recommendations in the code documentation of (5)). The resulting p-value distributions are shown in Figure 1B. Despite using a different test statistic and projection strategies, the same qualitative pattern is observed: p-values derived from the real dataset deviate strongly from those obtained from the randomized control, again producing significant rejections of the null hypothesis.

Finally, we implemented the full statistical test proposed by González-Delgado et al. (4), which computes p-values directly for Wasserstein-based statistics on the torus. As shown in Figure 1C, this method yields similar results: the randomized dataset produces the expected null distribution of p-values, while the real dataset displays an enrichment of low p-values leading to FDR-controlled rejections.

Taken together, these results demonstrate that the observed codon-conditioned differences in backbone dihedral angle distributions are not an artifact of a particular statistical test. Instead, the signal persists across multiple independent statistical frameworks and parameter choices.

Our reanalysis demonstrates that statistically detectable differences between synonymous-codon-conditioned backbone angle distributions remain observable when using statistically valid testing procedures and alternative test statistics, including those proposed by González-Delgado et al. While these results do not establish a mechanistic explanation for the observed signal, they indicate that the possibility of codon-dependent conformational effects cannot be ruled out on statistical grounds alone. Several experimental studies have shown that individual synonymous mutations can influence protein folding or function (7-10), suggesting that translation dynamics may in some cases influence structural outcomes. Whether such mechanisms contribute to the signal observed here remains an open question.

Further progress in addressing this question will likely require larger datasets that combine structural information with the exact coding sequences used to produce proteins. At present, such analyses are hindered by the fact that coding sequences are rarely deposited alongside protein structures in structural databases. In many structural studies, genes are codon-optimized or otherwise modified prior to expression, yet the resulting DNA sequences are typically not archived with the structural data. We therefore suggest that structural databases such as the Protein Data Bank consider enabling deposition of the coding sequences used in structural studies. Such information would facilitate systematic investigation of the relationship between genetic coding and protein structure.

## Methods

The distribution of p-values obtained when comparing Ramachandran angle distributions between pairs of synonymous codons was calculated using three different methods. Each method was run on the data in (1 and denoted real) and a randomized version thereof (denoted randomized). The randomized control dataset was created by randomly assigning codons to each amino acid according to empirical codon frequencies measured within the same secondary-structure class across the E. coli proteome.

The first method followed the procedure presented in (1) but with removal of the bootstrapping resampling procedure and standard permutations (K=5000). The results are shown in Figure 1A.

The second method followed the procedure in (1) but with removal of the bootstrapping resampling procedure and standard permutations (K=5000) and replacement of the KDE-L1 test statistic with Projected Wasserstein distance on the torus implemented as in (6) using with either 2 random or 4 fixed projections. The fixed projections were chosen as the recommendations in the code documentation of (5). The results are shown in Figure 1B.

The third method implemented the full statistical test proposed by González-Delgado et al. (4) and the results are shown in Figure 1C.

In all cases the Benjamini–Hochberg false-discovery-rate control (FDR) was set to 0.05 Code for reproducing our experiments is available at 10.5281/zenodo.16848517

